# Genomic epidemiology and evolution of *Escherichia coli* in wild animals

**DOI:** 10.1101/2020.07.23.216937

**Authors:** Robert Murphy, Martin Palm, Ville Mustonen, Jonas Warringer, Anne Farewell, Danesh Moradigaravand, Leopold Parts

## Abstract

*Escherichia coli* is a common bacterial species in the gastrointestinal tracts of warm-blooded animals and humans. Pathogenic and antimicrobial resistance in *E. coli* may emerge via host switching from animal reservoirs. Despite its potential clinical importance, knowledge of the population structure of commensal *E. coli* within wild hosts and the epidemiological links between *E. coli* in non-human hosts and *E. coli* in humans is still scarce. In this study, we analysed the whole genome sequencing data of a collection of 119 commensal *E. coli* recovered from the guts of 68 mammal and bird species in Mexico and Venezuela in the 1990s. We observed low concordance between the population structures of *E. coli* colonizing wild animals and the phylogeny, taxonomy and ecological and physiological attributes of the host species, with distantly related *E. coli* often colonizing the same or similar host species and distantly related host species often hosting closely related *E. coli*. We found no evidence for recent transmission of *E. coli* genomes from wild animals to either domesticated animals or humans. However, multiple livestock- and human-related virulence factor genes were present in *E. coli* of wild animals, including virulence factors characteristic for Shiga toxin-producing *E. coli* (STEC) and atypical enteropathogenic *E. coli* (aEPEC), where several isolates from wild hosts harboured the locus of enterocyte effacement (LEE) pathogenicity island. Moreover, *E. coli* in wild animal hosts often harboured known antibiotic resistance determinants, including against ciprofloxacin, aminoglycosides, tetracyclines and beta-lactams, with some determinants present in multiple, distantly related *E. coli* lineages colonizing very different host animals. We conclude that although the genome pools of *E. coli* colonizing wild animal and human gut are well separated, they share virulence and antibiotic resistance genes and *E. coli* underscoring that wild animals could serve as reservoirs for *E. coli* pathogenicity in human and livestock infections.

**Importance:** *Escherichia coli* is a clinically importance bacterial species implicated in human and livestock associated infections worldwide. The bacterium is known to reside in the guts of humans, livestock and wild animals. Although wild animals are recognized to serve as potential reservoirs for pathogenic *E. coli* strains, the knowledge of the population structure of *E. coli* in wild hosts is still scarce. In this study we used the fine resolution of whole genome sequencing to provide novel insights into the evolution of *E. coli* genomes within a broad range of wild animal species (including mammals and birds), the co-evolution of *E. coli* strains with their hosts and the genetics of pathogenicity of *E. coli* strains in wild hosts. Our results provide evidence for the clinical importance of wild animals as reservoirs for pathogenic strains and necessitate the inclusion of non-human hosts in the surveillance programs for *E. coli* infections.

## Introduction

*Escherichia coli* is a common Gram-negative commensal bacterium that resides in the intestines and faeces of warm-blooded animals as the dominant strain of the corresponding microbiomes (1). The commensal nature of *E. coli* may facilitate its dissemination across hosts, which provide the bacterium with a constant supply of nutrients and protection against environmental stresses (2,3). Pathogenic and Antibiotic Resistant (AMR) clones of *E. coli* have spread rapidly over recent years. Because half of the total natural *E. coli* population is estimated to inhabit environmental sites and a significant number of acute *E. coli* infections are known to have zoonotic origins (2), non-human hosts and settings are large potential reservoirs for pathogenic and AMR strains and genes (4).

Despite its likely importance for human health, the genetic diversity of commensal *E. coli* within wild hosts is still poorly studied, primarily due to the difficulty of recovering samples. Some studies have suggested *E. coli* and colonized hosts to co-evolve, such that the genomic characteristics of *E. coli* depend on the host species (5,6). While neutral evolutionary forces likely dictate most of the *E. coli* genetic diversity, feeding habits, diet and the microenvironments of the gastrointestinal tracts of hosts may constitute powerful selection pressures driving the phenotypic differentiation of commensal strains. These factors have led to the result that *E. coli* from wild animals often fall into other genetic and phenetic clades and phylogroups than isolates retrieved from humans (5–7).

Reports have described high multidrug resistance in some environmentally sourced *E. coli*, but strains found in wild animals generally display lower AMR than those found in livestock and non-animal environmental samples. The proximity of wildlife to human settlements seems to influence the AMR of gut microbiomes in wild hosts, due to antibiotic pollution (8,9). Whether wild animals predominantly act as sources or sinks in AMR evolution is still unclear (10,11). A recent study in Nairobi found interactions between humans and livestock to catalyse the colonization of wildlife by AMR *E. coli*. Similar to AMR genes, the distribution of virulence factor genes within environmental *E. coli* isolates is still an understudied area, although accumulating evidence suggests that there are epidemiological links between pathogenic strains of *E. coli* in livestock and those in humans (12,13).

The poor resolution of typing methods, the limited diversity of host species under study and the sampling bias towards food animals in previous studies have substantially limited our understanding of commensal *E. coli* in wild hosts. Furthermore, the effects of human interventions in the habitats, in particular the exposure of wild hosts to mass-produced antimicrobials, complicate efforts to study the genetic determinants of commensalism. To address these limitations, we examined whole genome sequences of 119 commensal *E. coli* isolated recovered from the faecal samples of 68 wild mammals and birds host species from North America, predominantly from Mexico (5). With an estimated 2000 different resident species, Mexico hosts 10–12% of the world’s biodiversity (14), which offers the opportunity to scrutinize the host-pathogen evolution for a wide range of wild host populations.

Our results indicate that *E. coli* populations in wild hosts are only weakly associated with the taxonomy and ecological and physiological attributes of the host species. Furthermore, while we detected no recent epidemiological links with human strains, we observed local population mixing and some sharing of antibiotic resistant genes and virulence genes between strains from wild and domesticated/livestock animal hosts. These results highlight the subtle distinction between virulence and commensalism and implicate wild hosts as reservoirs for *E. coli* pathogens.

## Material & Methods

### Strain acquisition, sequencing and genome assembly

We acquired a systematic collection of commensal *E. coli* from a previous study (5). The collection comprised 119 faecal strains from hosts belonging to 81 animal species, 31 families and 16 orders. 110 and 9 strains were from mammals and birds, respectively. 110 strains were recovered from Mexico and the rest were isolated in Venezuela and Costa Rica during the 1990s. The antimicrobial susceptibility testing was conducted on the whole collection for 8 antimicrobials clinically approved for treating *E. coli* infections, including beta-lactams (ampicillin, cefotaxime, ceftazidime, cefuroxime and cephalothin), aminoglycosides (gentamicin and tobramycin), ciprofloxacin and trimethoprim, as described in (15). The full description of the strains with metadata is available in Supplementary Table S1.

DNA was extracted with the QIAxtractor (Qiagen) kit according to the manufacturer’s instructions. We prepared Illumina sequencing libraries with a 450-bp insert size and performed sequencing on an Illumina HiSeq2000 sequencing machine with pair-end reads of length 100 bp. Ninety-six samples were multiplexed to yield an average depth of coverage of ∼85 fold. Short reads data were submitted to the European Nucleotide Archive under the study accession number of PRJEB23294. Reads were then assembled and improved with an automated pipeline, based on Velvet with default parameters. Assemblies were annotated with an improvement assembly and Prokka-based annotation pipeline, respectively (16–18). Details on assembly statistics and gene annotations are available in Supplemental Table S1. Roary, with the sequence identity value of 95% for orthologous groups, was used to create a pan-genome from annotated contigs (19).

Multilocus sequence typing was performed on assemblies using a publicly accessible typing tool anddatabase,availableon(www.github.com/sanger-pathogens/mlst_check)withdefault parameter values to identify ST clones. We contextualized our collection with *E. coli* strains from environment, livestock/domesticated animals and humans in the publicly available Enterobase dataset (www.enterobase.warwick.ac.uk). Since we were primarily interested in recent evolution and transmissions between *E. coli* in wild hosts and other hosts, we retrieved genomic data and metadata for all strains with an identical ST with at least one strain in our collection on 26/04/2020. We included only strains for which prior consent was obtained from the strain’s owners. In total, genomic data for 1826 strains was retrieved. We then classified strains based on their source of isolation as environment, livestock/domesticated animals and human associated. We used the above-mentioned pipeline to assemble the pair-ended short reads and annotate the assemblies also for external samples.

### Mapping, variant calling and phylogenetic analysis

We mapped short-read sequences to the *E. coli* K12 sequence (Biosample id: SAMN02604091), with SMALT v 0.7.4 (www.sanger.ac.uk/resources/software/smalt/), with a minimum score of 30 for mapping. SAMtools and BCFtools were then employed to annotate SNPs (20). SNPs at sites in which SNPs were present in less than 75% of reads were excluded. We extracted SNPs from the core-genome alignment produced by Roary and mapping to the *E. coli* K12 reference genome using the script available at https://github.com/sanger-pathogens/snp-sites.

To construct the alignment-free phylogenetic trees, we first enumerated *k*-mers of size 50 from assemblies with the frequency-based substring mining(fsm-lite)package (www.github.com/nvalimak/fsm-lite). We subsequently counted the number of identical *k*-mers for pairs of isolates to produce a similarity matrix, which was then converted into a distance matrix. The distance matrix was used as input for the ape (21) and phangorn (22) packages to produce a neighbour-joining phylogenetic tree. The tree was visualized with iTOL (23) and Figtree (www.tree.bio.ed.ac.uk/software/figtree/).

### Virulence factors, antimicrobial resistance genes identification and *in silico* serotyping and LEE typing

Virulence factors and antimicrobial resistance genes were identified with the srst2 (24) package using the Virulence Factor Data Base (VFDB) (25) and ResFinder database (26) available in the package, respectively. We employed a loose similarity cut-off of 60% to ensure that divergent genes were detected.

The genomic context of the AMR genes was explored in two ways. First, we searched the Nucleotide database to find similar annotated genomic regions with the contig that contain the resistance gene with blastn. Second, to further examine whether genes are located on plasmid or chromosome, we also utilized PlasmidSPAdes (27) to first reconstruct plasmid assemblies and then screened the contigs for the AMR gene with blast, as part of the assembly graph viewer Bandage (28). We identified LEE loci and serotypes with the typing method in the srst2 package, using a similarity threshold of 60%. We then confirmed the presence of virulence factor genes by running blastn against assemblies. For the O-antigens produced by Wzy-dependent pathway, variations in unique genes *wzx* (encoding an O-antigen flippase) and *wzy* (encoding an O-antigen polymerase) were examined (29). For the ABC transporter-dependent pathway, variations in *wzm* (encoding an O-antigen ABC transporter permease gene) and encoding *wzt* (encoding an ABC transporter ATP-binding gene), involved in O-antigen synthesis, were studied.

### Association with ecological and taxonomical attributes of host species

We obtained the tree of life for the host wild host species with the R package rotl (30) and visualized the concordance between the host tree and the core genome tree of colonizing *E. coli* strains with Dendroscope (31). We used treedist function in ape package to compute the distance matrix from phylogenetic tree. For *E. coli* strains, the distance matrix was obtained from pairwise Hamming distances between core genome sequences. We then used a Mantel test with 1000 permutations as part of ade4 package (32) to assess the correlation between the distance matrices for *E. coli* genomes and that for host species. To compute the difference between the phylogenetic trees of *E. coli* strains and hosts, we used the treedist function, as part of the phangorn package in R. By doing so, we computed the square root of the sum of squares of differences in path length between each pair of tips in two trees (33). The path is defined as the number of edges within the tree that must be traversed to navigate from one tip to the other.

Furthermore, we dissected the relationship between virulence ability, measured as the total number of virulence genes, and ecological and physiological attributes of each host species in the panTHERIA database (34). The database includes a comprehensive species-level data set of life-history, ecological and geographical traits of all known extant mammals. Spearman’s rank correlation coefficient values were computed to assess the significance of the correlation between virulence gene count and attributes.

### Positive selection analysis

We analysed positive selection by reconstructing the ancestral sequence for each gene in the core genome, identified by Roary, with FastML (35). Subsequently the seqinR 1.0-2 package (36) was employed to compute the *K*_*a*_ and *K*_*s*_ values for each strain, in comparison to the ancestral sequence. We left out the strains with no synonymous changes, i.e. *K*_*s*_==0. For functional enrichment analysis, COG categories of genes were extracted from the annotation by Prokka and assigned to functional classes.

### Bayesian analysis

We constructed a Bayesian tree using the BEAST (37) to date the recent mixing between *E. coli* from wild hosts and other strains in a clone in the B1 phylogroup. The clone was identified with the clustering tool in adegenet package (38). To this end, we screened the SNP cut-off value for identifying clusters in the wild host and global collection and used the clustering that remained unchanged for the highest number of SNP cut-off values. We then extracted the cluster that contained a high number, i.e. 39/119, of *E. coli* strains from wild hosts.

We mapped the short reads for strains in the cluster to a local reference genome, i.e. the strain with the lowest number of contigs. We then ran Gubbins (39) with 5 iterations to remove hypervariable sites from the genome alignment and produced a neighbour-joining phylogenetic tree. To assess the strength of the temporal signal, we plotted the root-to-tip distance versus year of isolation and performed 10,000 bootstraps with randomized years to attain a distribution for R-squared values. Subsequently, we compared the R-squared value for the data distribution with the simulated distribution. The temporal signal was 40% confidence for the clone under study.

The multiple alignment was then used as input for BEAST. We examined a range of prior models, including a strict molecular clock and a log-normal model of a relaxed molecular clock with constant population size. Markov chain Monte Carlo (MCMC) simulations were performed three times for 50 million generations with sampling every 10 generations. A cut-off of 200 was chosen for the Effective Sample Size (ESS) of key parameters for the convergence. The 95% Highest Posterior Interval (HPI) was used to report the certainty on ages of ancestral nodes.

## Results

We sequenced 119 strains from 68 wild animal host species and found them to capture much of the known *E. coli* genetic diversity. Indeed, our wild host population contained representatives of all of the major known phylogroups of *E. coli*, with group B1 (55 strains, 47% of all) being most prevalent followed by B2 (21 strains, 18%), A(17 strains, 14%), D (15 strains, 13%) and E (7 strains, 6%) (Figure 1A). The high frequency of B1 strains is consistent with previous epidemiological reports on *E. coli* isolated from domesticated animals but stands in contrast to the high prevalence of phylogroups B2 and A among *E. coli* isolated from human (40). *E. coli* from domesticated/livestock animals and North America were disproportionately likely to share phylogenetic origin with our wild *E. coli* strains (Figure S1A, S1B), suggesting regional dissemination of some *E. coli* phylogroups across both domesticated/livestock and wild animals.

**Figure 1.**
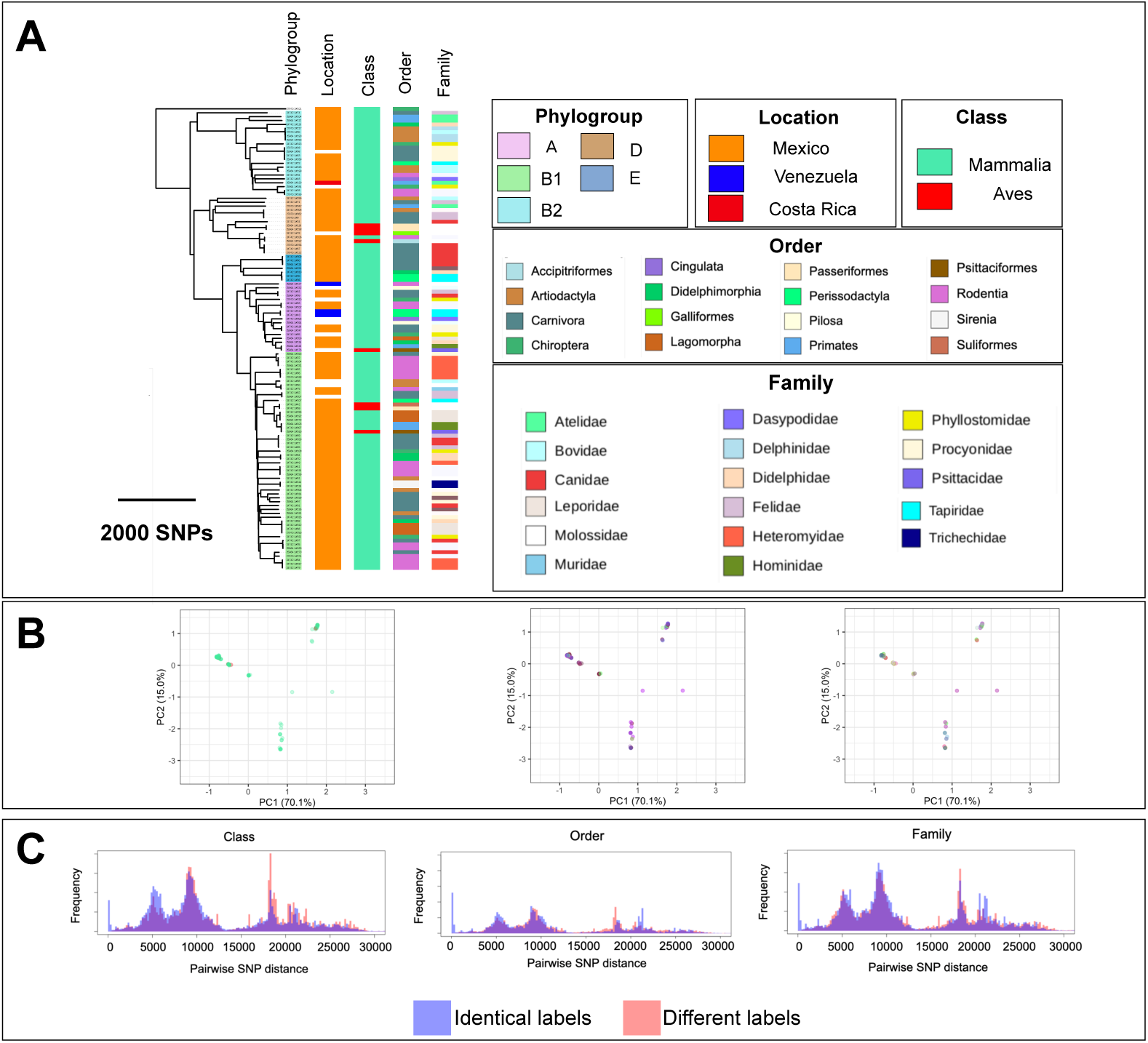
Phylogenetic distribution of host specificity and cluster analysis: A) phylogenetic tree of 119 *E*.*coli* strains from wild hosts and its association with host taxonomic level. Families represented by one strain are not shown. B) Principal component analysis of the strains, with labels of the phylogroup and taxonomic rank. Each colour corresponds to one taxonomic rank. Families represented by one strain are not shown. C) Distribution of pairwise SNP distances for strains belonging to the same (red) and different (blue) taxonomic rank.

The concordance between the evolutionary histories of *E. coli* and their hosts was significant. Both comparisons of host and *E. coli* distance matrices (*p*=0.0001, Mantel test, Figure 2A) and distances between phylogenetic trees for *E. coli* strains and hosts to distances in randomized trees (*p* = 0.003, 1000 tests, Figure 2B) rejected completely random observations. Despite this, we found only a moderate correlation of 0.47 between the genetic distance matrices for *E. coli* strains and hosts (Figure 2A), with closely related *E. coli* often colonizing divergent wild hosts and closely related wild animal species often hosting distantly related *E. coli*. The weak genetic association between *E. coli* and their wild hosts is also evident at higher taxonomic levels, with only weak genetic clustering of *E. coli* according to the host class, order and family (Figure 1B). This is further confirmed by the extensive overlap in the distributions of SNP distances for *E. coli* pairs colonizing host species from the same taxonomic groups and those of pairs colonizing different taxonomic groups (Figure 1C), as 0.95, 0.95 and 0.96 of ranges of distributions overlapped for taxonomic ranks of class, order or family, respectively. The accessory genome of *E. coli* colonizing wild hosts has evolved in concert with the core genome (*p*=0.0001; Pearson’s R=0.85, Mantel test on distance matrices for core genome and accessory genes; Figure S2). Thus, we found little evidence of horizontal gene transfer between lineages colonizing wild animals.

**Figure 2.**
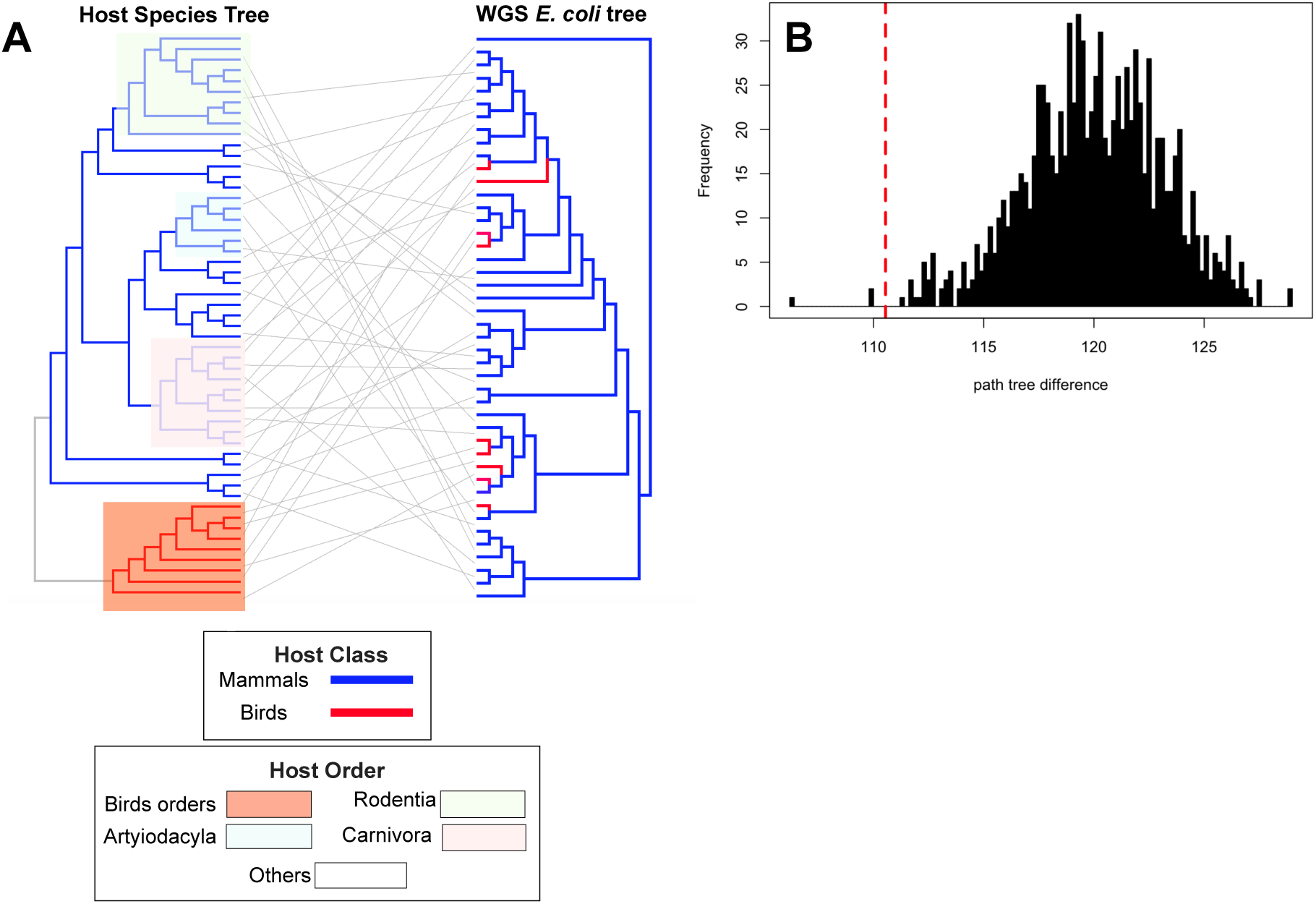
Concordance between host and *E. coli* phylogenetic trees: A) Phylogenetic tree of the whole genome sequencing of *E. coli* strains and the Tree of Life (TOL) for host strains. For host species for which more than one isolate was available in the dataset, one strain was randomly drawn. Clades for bird and major mammalian orders are highlighted. B) The distance, i.e. the path tree difference, between the trees in A) shown in dotted red line. The black bars are the distribution of path tree differences, computed for 1000 trees that were generated by randomly shuffling tree tips of the host tree in A).

We compared the rates of non-synonymous and synonymous single nucleotide evolution (*K*_*a*_/*K*_*s*_) since the last shared common ancestor of *E. coli* colonizing wild animals. Out of 3,659 genes in the core genome, 253 genes had at least one *K*_*a*_/*K*_*s*_ value above 1, with an average of 11.7 genes, i.e. 0.3% of total genes, per strain falling in this category (Figure S3A, S3B). The number of genes under strong positive selection did not show any link with host (Figure S3B). The strongly selected genes encoded proteins involved in a broad range of functions, including energy production, carbohydrate and ion metabolic and transport and signal transduction proteins being slightly overrepresented (Figure S3B). Thus, many diverse functions may have been involved in adapting *E. coli* to commensalism in different wild animals and genome-wide selection have not been affected by host species.

We next probed the genomic evolution and epidemiology of *E. coli* colonizing wild-animals in relation to those of the global *E. coli* collection coming from other hosts. We found no evidence for recent *E. coli* transmission from wild animals to human hosts or domesticated/livestock hosts, underscoring the role that ecological and geographical barriers played in limiting *E. coli* spread (Figure S1A). The most closely related *E. coli* strains colonizing wild animals and humans were separated by 40 SNPs in their core-genomes, which, assuming a substitution rate of two SNPs/year (41), corresponds to 20 years. However, in a number of cases, we found signs of *E. coli* colonizing wild animals to have diverged recently from *E. coli* colonizing domesticated animals. This was particularly evident in the B1 phylogroup where one-third of our *E. coli* from wild animals, clustered with lineages isolated from domesticated/livestock animals (*n* = 96), food (*n* = 12) and environmental sources (*n* = 13). We reconstructed the Bayesian tree of these 158 strains and found their last common ancestor to have lived about 1000 years ago, with a substantial expansion of the clade over the past 100 years (Figure 3). We identified eight incidents of strains jumping between wild animals and other sources in this clade, all during the last 100 years and all but one during the last 50 years (Figure 3). One recent incident involved *E. coli* jumping between wild hosts residing in city regions and domesticated/livestock animals. These *E. coli* host switching events may reflect anthropogenic intervention in the habitats of wild hosts, and the rapid urban and agricultural growth and environmental degradation in Mexico over the past decades (42).

**Figure 3.**
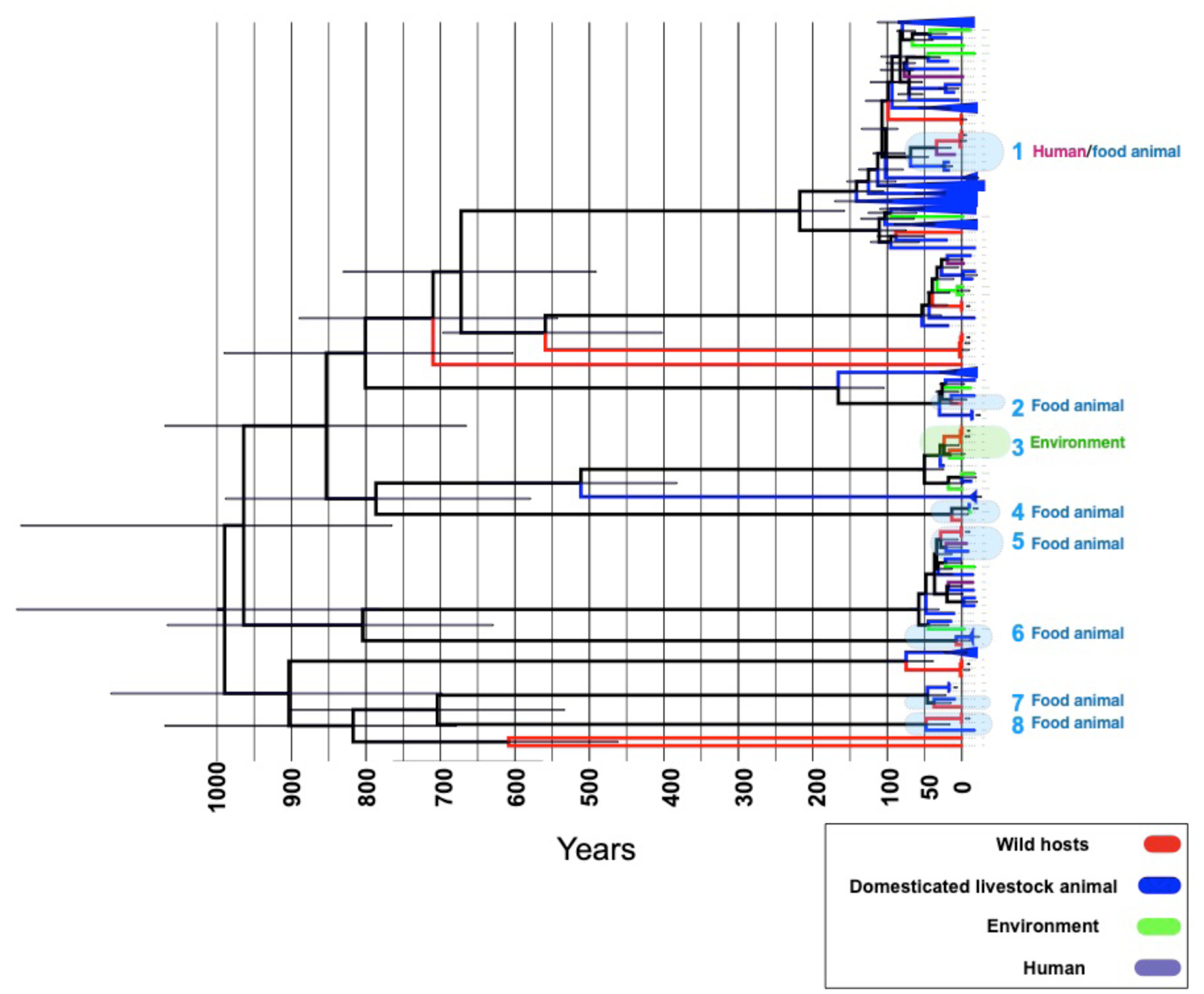
Recent mixing of wild and non-wild hosts lineages: Bayesian tree for strains in a clade belonging to B1 phylogroup. The shaded boxes show putative host jumps events between wild hosts and other sites, i.e. domesticated animals, environment and humans, over the past 100 years. The error bar shows the 95% confidence interval from the Bayesian tree.

The recent *E. coli* jumps between wild and domesticated animals led us to examine whether *E. coli* colonizing the former harbour any known human or food-animal-linked virulence factors. We identified a range of virulence factor genes, including four types of toxin genes, two adhesin genes, two iron chelators and three transporters. These were present in *E. coli* colonizing different wild animals (Figure 4A). The frequency of virulence factors was on average higher for strains recovered from Primate (11.5 genes per isolate), Rodentia (9.5 genes per isolate) and Carnivora (12.5 genes per isolate) host species (Figure 4A, 4B). Some species not closely related to humans, such as birds, were colonized by strains carrying a high number of virulence factors (Figure 4A, 4B), suggesting that the pattern is not a reflection of the higher frequency of human- and livestock-associated genes in the database.

**Figure 4.**
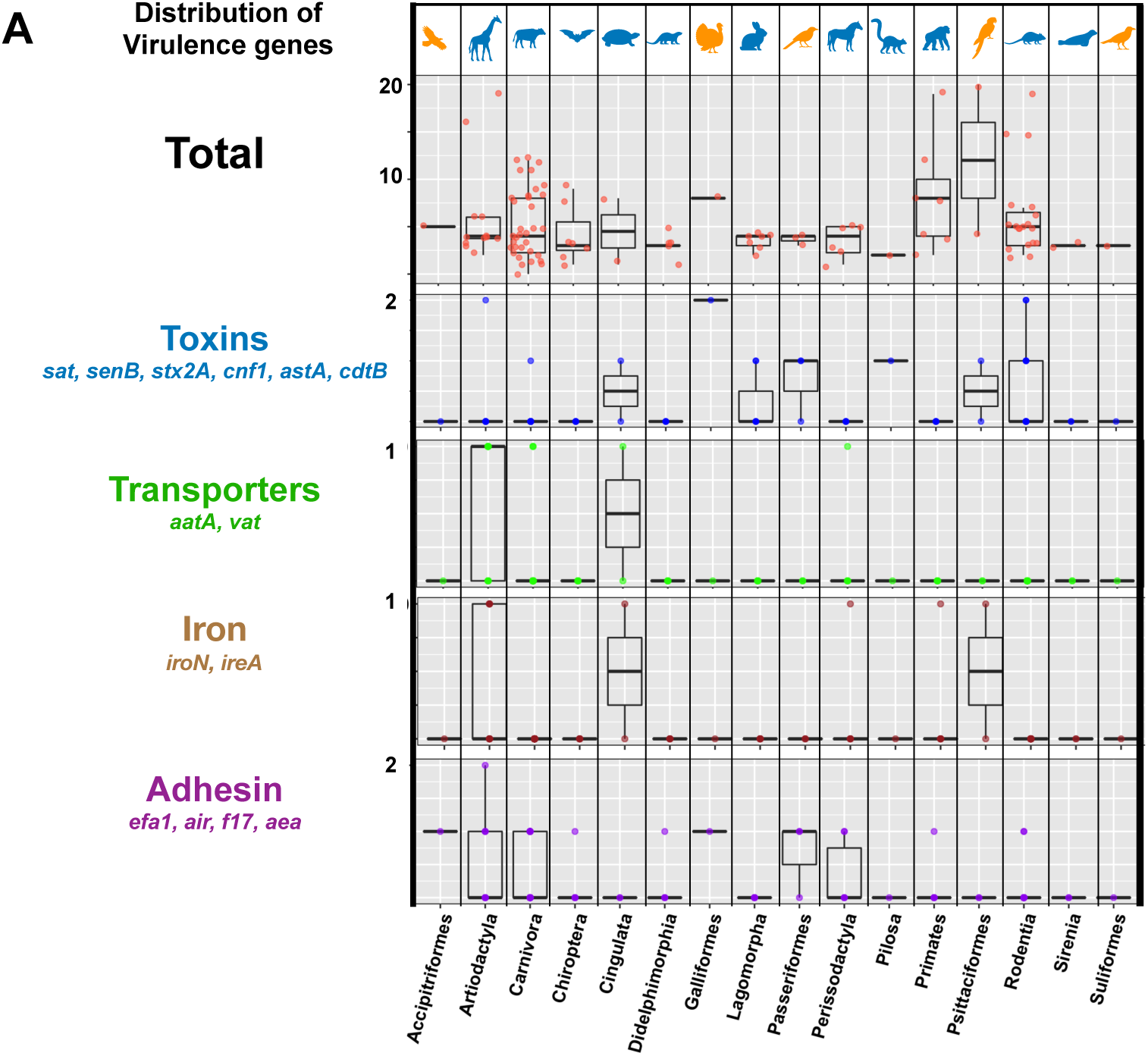
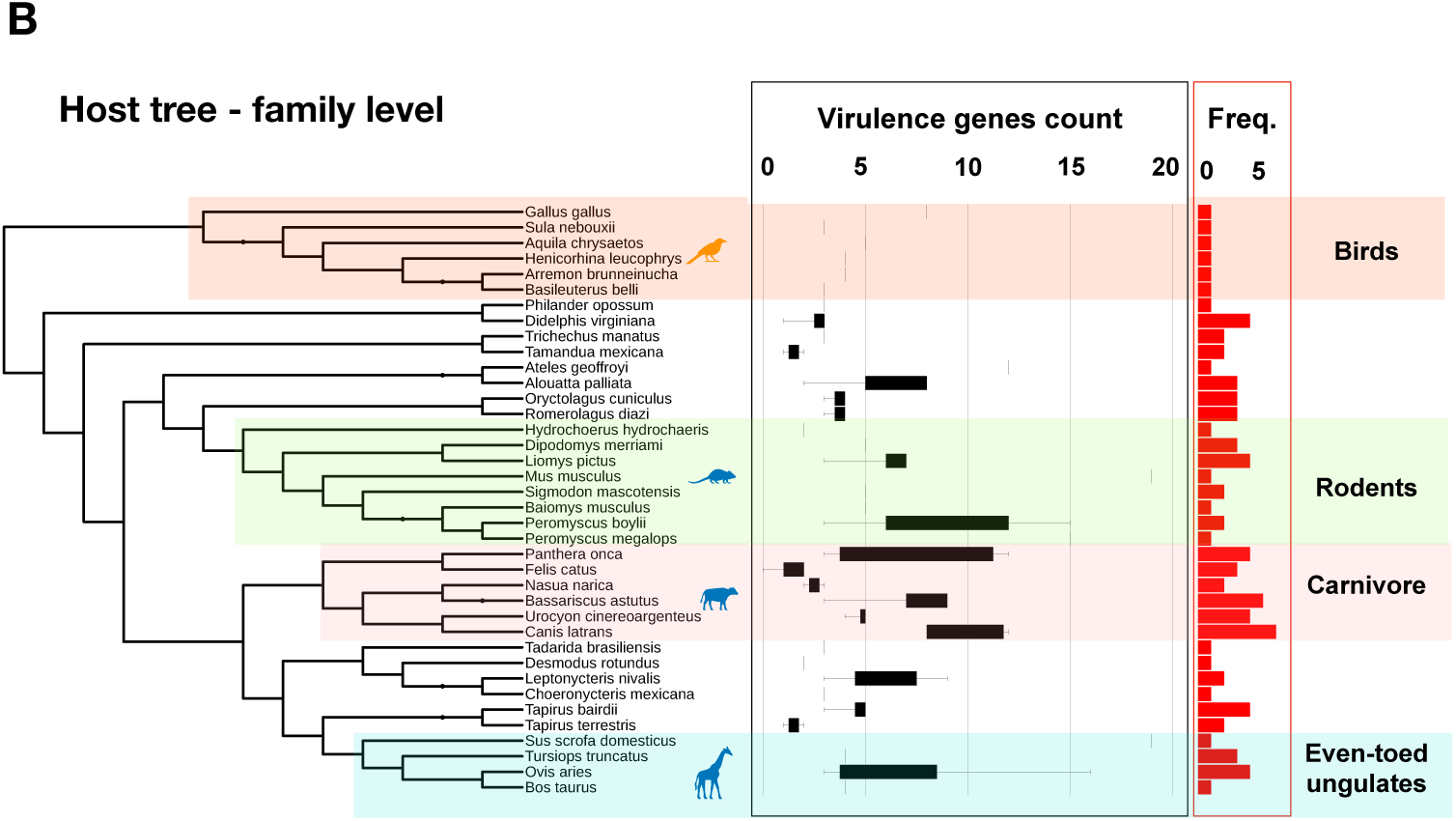
Distribution of virulence factor genes: A) The frequency of virulence factors genes across functional groups and taxonomic orders. B) The phylogenetic distribution of *E. coli* virulence genes across wild animal host species. The tree shows the tree of life for hosts, where major orders are shown in shaded boxes. Bar plots show the frequency of genes. Horizontal boxplots represent the distribution of virulence genes for strains recovered from each host across host orders.

Because both the physiology and ecology of the host species can affect the virulence factors encoded in the genomes of infectious bacteria, we examined the relationship between the number of virulence genes in *E. coli* colonizing wild animals and the 45 such features in the panTHERIA database. A previous study on four virulence genes revealed that body mass of the host species is positively linked with the number of virulence factors present in the gut microbiome and this was attributed to the gut complexity (43). However, our analysis on more virulence genes showed no such correlation, considering either adult, neonate or weaning body mass (Figure S4A). Only habitat breadth (*p*=0.013, Spearman’s *ρ*=-0.23), diet breadth (*p*=0.015, Spearman’s *ρ*= -0.26) and social group size (*p*=0.002, Spearman’s *ρ*=0.29) correlated significantly with virulence gene counts, with more diverse habitats and diets associating to fewer, and formation of larger social groups to more, virulence genes (Figure S4A, S4B). Larger social groups, as observed in Carnivora, Artiodactyla and Primates in Figure S4C, is known to increase the social transmissions of infectious agents in animal societies, which may facilitate the dispersion of virulence genes (44). Although a larger sample set is needed to examine the impact of potential confounding factors, the findings further support the idea that a complex network of host- and the environment-related factors shapes the genomic characteristics of commensal strains.

Certain *E. coli* serotypes, which reflect O, H and K antigen variation and not necessarily evolutionary relatedness, are recognized to cause virulence in human and livestock associated-infection. We found 53 and 14 serotypes to be shared between *E. coli* strains in wild hosts and those in domestic animal and human infections, respectively (Supplemental Table S1). In total, we identified 71 distinct serogroups and 14 strains that were not typeable among *E. coli* colonizing wild animals, further underscoring their broad diversity. The serogroups of 74 strains overlapped with those of known pathovars, including non-O157 Shiga toxin-producing *E. coli* (STEC) (*n* = 40), enterotoxigenic (ETEC) strains (*n* = 12), enteropathogenic (EPEC) strains (*n* = 11) and enteroaggregative (EAEC) strains (*n* = 11), across hosts (Figure 5A) (Supplemental Table S1). The pathovars are recognized to have non-human sources and may be acquired via direct contact with either animals or their faeces in petting zoos and on farms (for STEC) or through the consumption of contaminated water and food (for EAEC and ETEC), as previously reported in Mexico (32,33). ETEC is also an important cause of diarrhoea in domestic animals, notably calves and piglets (45). Two strains from the wild hosts collection shared serotypes with pathogenic strains and contained genetic virulence hallmarks of their associated pathovars. One strain belonged to O111:H8, a clinically relevant enterohemorrhagic *E. coli* (EHEC) serotype, and contained both the enterocyte effacement (LEE) pathogenicity island (PAI) and the toxin *stx2* gene. This strain was recovered from a wild sheep close to a city. The other strain belonged to O78:H34 and was isolated from a parakeet carrying enteroaggregative *E. coli* (EAEC) virulence genes, including the plasmid-encoded, heat-stable enterotoxin toxin (EAST-1) and *aatA* and *aggR*, encoding a transporter of a virulence protein and a virulence regulator, respectively (Figure 3A). The serogroup was recently isolated from free pigeons in Brazil, showing the circulation of the pathovar amongst birds (46). None of the serotypes associated with STEC and ETEC pathovars were found to carry a toxin gene. Although the sharing of serotypes with pathovars does not necessarily cause the strain to become virulent, our serotype analysis further underscored the genetic overlap between *E. coli* in wild and food animals (see the discussion section).

**Figure 5.**
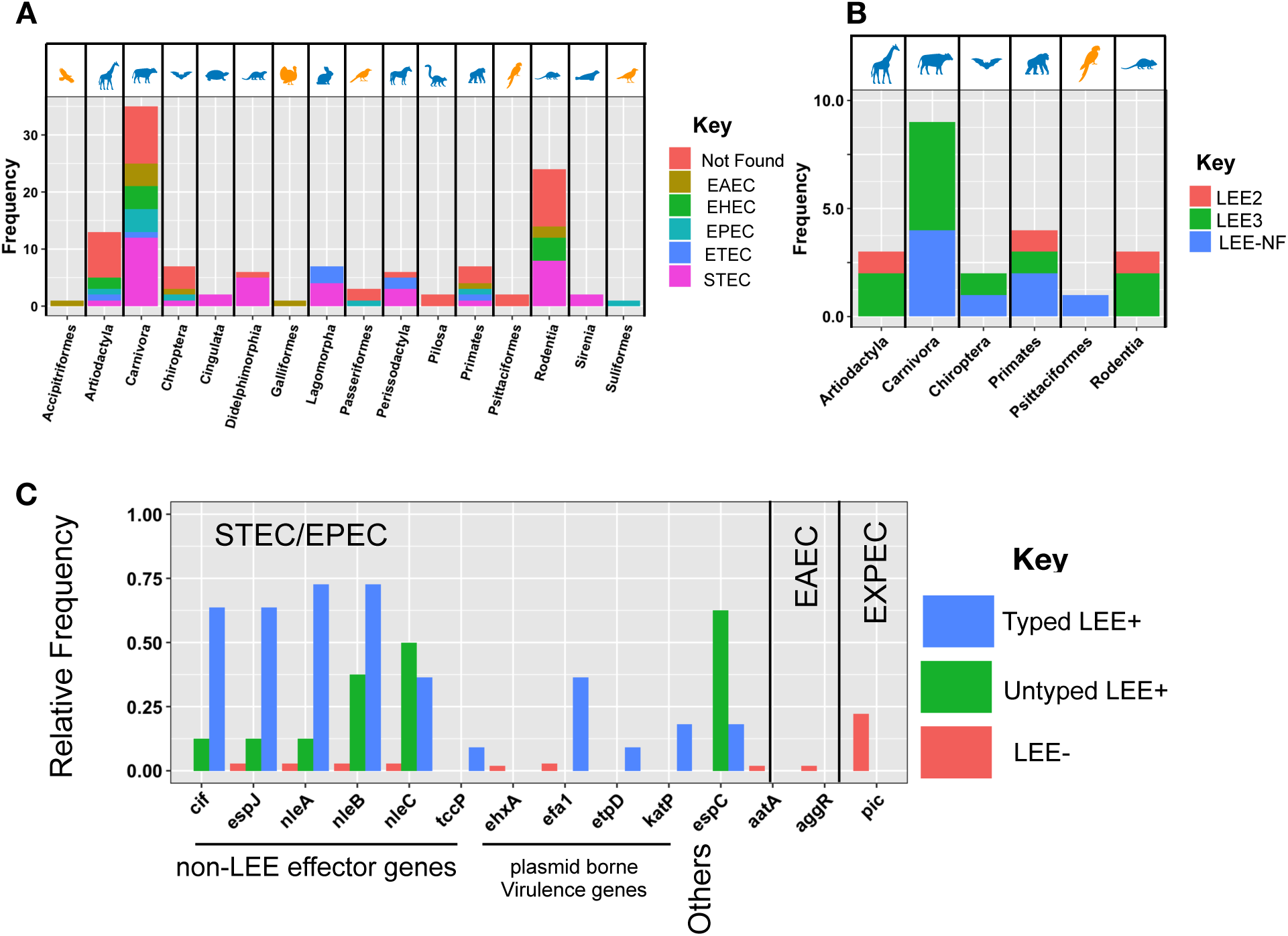
Sharing of serotypes and distribution of LEE genes and effectors genes across hosts: A) Distribution of serotypes shared between *E. coli* colonizing wild hosts and known pathovars across taxonomic orders of hosts. B) Distribution of typed and non-typed LEE families across taxonomic orders of hosts. B) Distribution of virulence genes and LEE effector genes in typed, untyped LEE(+) and LEE(-) strains.

We found the enterocyte effacement (LEE) pathogenicity island locus, a hallmark of STEC and EPEC pathovars, in 21 of the *E. coli* lineages from wild animal hosts and these hosts belonged to six different taxonomic orders (Figure 5B, Supplemental Table S1). The locus encodes factors required for the colonization of the human intestine (47). However, the absence of the plasmid carrying *E. coli* adherence factors (pEAF) led us to classify these isolates as atypical EPEC (aEPEC), an *E. coli* class widely spread across food animals and humans (48). The LEE-positive strains also harboured other virulence factors that are typical of EAEC and EXPEC pathovars and affect pathogenicity (Figure 5C). This included genes normally located on STEC virulence plasmids, such as pO157, pO26, *espP* and *nle*, all of which were significantly more frequent in LEE-positive strains than in LEE-negative strains (Figure 5C). We found 2 and 11 strains, all in the B1, E and D phylogroups, to carry the LEE2 and LEE3 variants respectively, while 8 strains, mainly in the B2 phylogroup, carried a non-typeable LEE loci. All three loci types were broadly distributed among host taxonomic families, in agreement with them benefitting *E. coli* colonization of animal guts in a general sense, as previously proposed for bovine hosts (12). Our findings also agree with the virulence ability of aEPEC strains spanning across a broad host range of with that virulence in STEC and EPEC strains evolved by commensal strains acquiring virulence factors sequentially (48).

We found the *E. coli* collection in wild hosts turned to be sensitive to most antibiotics, except for ampicillin, against which 65% of strains were resistant (Supplemental Table S1). Their general susceptibility indicates the lack of exposure of wild animals to therapeutic levels of antimicrobials. Despite this, a range of AMR genes against beta-lactamase, aminoglycosides, sulfonates and ciprofloxacin were identified across different lineages and hosts species (Figures S5). The discordance between AMR phenotypes and genotypes points to regulation mechanisms or epistatic interactions, affecting the penetrance of resistance genes. The genomic context of AMR genes turned out to be diverse, with genetic linkage to a range of phage genes and IS elements, including to IS91 and IS10. For AMR genes located on sufficiently long contigs, we explored the genomic context and found similarity with a broad host range Col-plasmids (*n* = 21) and chromosomal (*n* = 3) regions. The genomic contexts varied across host species; for example, while one strain from a *Pilosa*, a placental mammal, harboured an AMR gene cassette consisting of *tet, str* and *sul* genes, for four strains from different mammalian species, AMR genes were found sporadically distributed across the genome. Besides plasmid-borne resistance determinants, we identified a set of ciprofloxacin-resistance mutations in the *parE*, and *gyrA* genes, independently emerged across lineages (Figures S5). The strains were recovered from Carnivora, Rodentia and Passeriformes species. Four of the isolates belonged to the clinically relevant O17/77:H18 serotype, which forms a highly relevant pathogenic group in the phylogroup D. This group of *E. coli* were an emergent clinical threat in the 1990s, predominantly in North America (49). Ciprofloxacin was introduced into clinical settings in the 1980s (50), prior to the sampling time period of our collection. The presence of ciprofloxacin resistance determinants in wild hosts, therefore, suggests that either rapidly emerging resistance was transmitted from wild hosts into human settings prior to the sampling time period, or the pre-existence of resistance in wild hosts reservoirs.

## Discussion

We provided insights into the evolution of the genetic repertoire for commensal lifestyles in wild hosts. The genome of wild host *E. coli* was stable and evolved mostly independently from host species. Certain lineages were recently mixed with *E. coli* strains from local domesticated/companion animals. Moreover, some strains harboured virulence and AMR genes shared with pathogenic human and livestock animal strains, highlighting the subtle distinction between pathogenicity and commensalism in *E. coli*. We note that since our strains were recovered from faeces, we are unable to delineate between pathogenicity and commensalism for our strains and to ascertain whether strains cause virulence when introduced into the blood stream.

The ability of diverse strains to colonize similar host species and the diverse range of virulence factors in *E. coli* from wild hosts point to the flexibility of the *E. coli* genome. These factors provide a flexible genomic repertoire for adapting to diverse host environments, consistent with the coincidental hypothesis of virulence (1,51,52). The evolution of virulence is complex and driven by opposing forces. Higher virulence leads to higher survival within the microenvironment of a specific host’s intestine but may harm the host, thus constraining the host range. Furthermore, the virulence factor gene may entail a fitness cost, leading to a loss of virulence in the long term. Here, our results provide evidence in favour of reduced host specialism, suggesting that a high level of versatility allows better domination and exploitation of resources in the evolution of *E. coli* (53,54).

The absence of recent divergence incidents between *E. coli* isolated from human and wild animals host suggests a clear separation between these groups. Hence, wild hosts are unlikely to have served as sources of recent human clinical infections. Such a clear genetic distinction between *E. coli* lineages in non-human and human settings has been suggested by several recent genomic epidemiological studies (55–57). However, *E. coli* colonizing in wild animal hosts may still serve as reservoirs for individual virulence or AMR genes, which can be transferred to pathogenic strains through HGT, or as genomic backbones which upon acquisition of further virulence factors may evolve into pathogens that can jump into human hosts. A recent large-scale genomic study has shown that livestock serve as an evolutionary source for human EPEC strains (12).

Our collection was predominantly recovered from Mexico in the 1990s. This clearly limits the scope of the implications of our results. In particular, over the past three decades, the rapid consumption of antibiotics, globalization, anthropogenic interventions in the wild, and contaminations of environmental sources potentially selected for higher virulence and resistance and facilitated jumping between human-wild hosts. Another limitation of our study is that we did not examine the intra-host diversity of *E. coli* strains. Genetically distinct strains reside within the gut, and their genetic composition varies across the different regions of the gut. Although it is known that one or two resident *E. coli* clones most often dominate the microbial community in the gut (58) and is likely to be one of the strains recovered from each species in our study, a deeper sampling from each host is required to examine the effect of interactions between complex intestinal microbiota and *E. coli* in within-host adaptation. Such sampling would also allow an examination of whether virulence genes in the dominant clone confers any fitness advantage over other clones. Our study also neglected the differential expression of virulence genes in commensal strains (59), which determines the regulation and functional level of these genes. Therefore, the integration of transcriptomic, (meta-)genomic and metabolomic data in a follow-up study would complement our findings.

Studies on *E. coli* genomics are biased towards the characterization of pathogenic clinical strains under therapeutic conditions. Deciphering the genetics of commensalism is necessary for understanding the transition from commensalism to pathogenicity. Besides providing epidemiological insights, such knowledge informs us about new host-pathogen interactions that could be targeted in treating *E. coli* infections.

## Acknowledgment

We thank Jan Michels for providing the strains. This work was partially funded by a grant from the Centre for Antibiotic Resistance Research (CARe) at the University of Gothenburg to AF and grant number 2016-06503 from the Joint Programming Initiative on Antimicrobial Resistance (JPIAMR) to JW and AF. This work was in part supported by the Academy of Finland (grant 313270 to VM). LP was supported by Wellcome, and Estonian Research Council (IUT34-4). DM was supported by the Joint Programming Initiative on Antimicrobial Resistance (JPIAMR) via MRC grant MR/R004501/1. The funders had no role in study design, data collection and analysis, decision to publish, or preparation of the manuscript.

## Supplemental Data

**Supplementary Figures Files**

**Supplemental Tables**

Supplemental Table S1: Samples specification, serotypes, associated pathovars and virulence and AMR genes

