## Supplemental Figures for "Genomic epidemiology and evolution of *Escherichia coli* in wild animals"

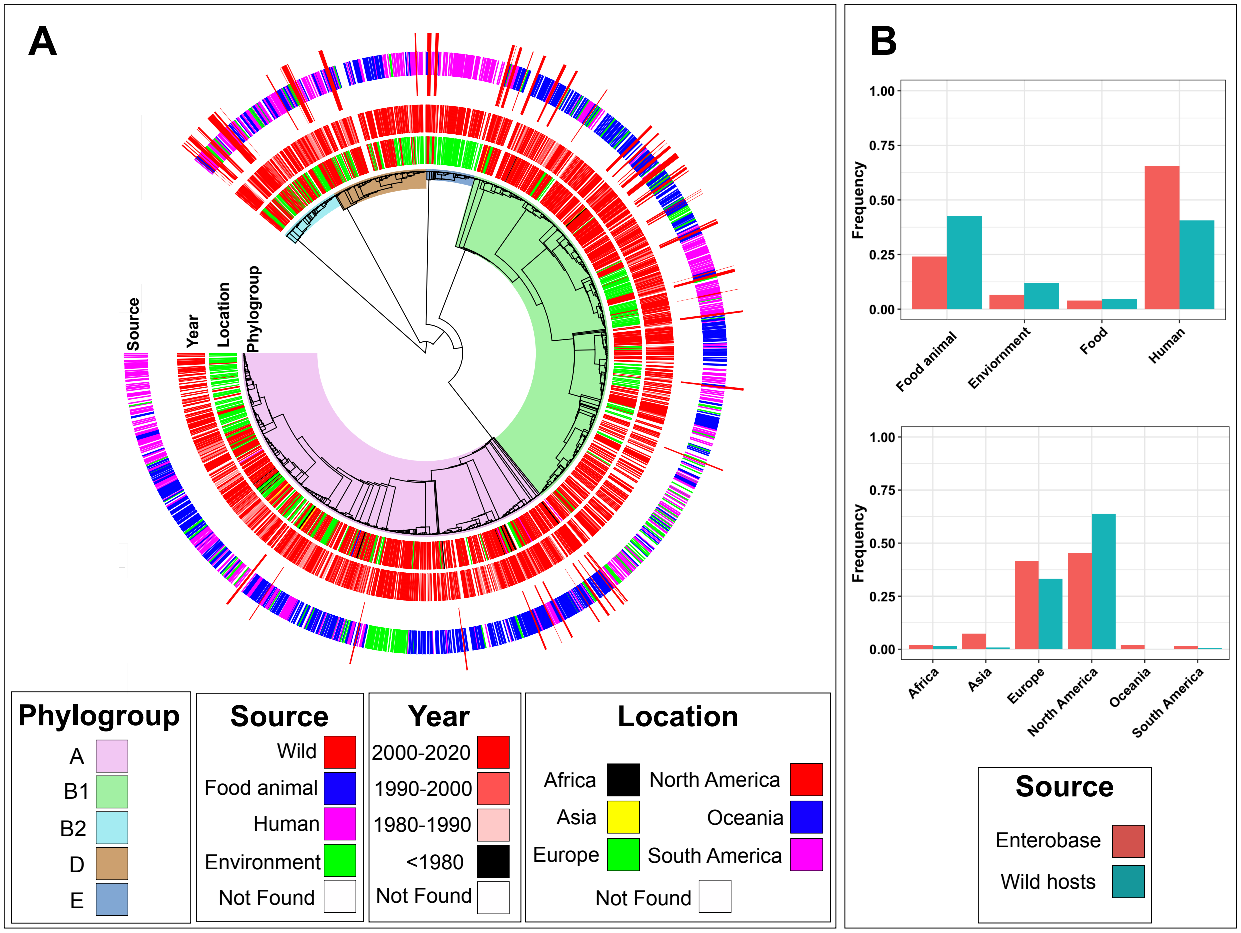


**Figure S1 Comparing the phylogeny of *E. coli* from wild hosts with strains from other sources**

A) Alignment-free phylogenetic tree for the *E. coli* strains in the context of previously sequenced genomes in the Enterobase database and the associated metadata of phylogroup and isolation sites. B-C) The distributions of continent of isolation and host in the curated dataset, compared with those in the Enterobase collection.

Robert Murphy, Martin Palms, Ville Mustonen, Jonas Warringer, Anne Farewell, Danesh Moradigaravand, Leopold Parts

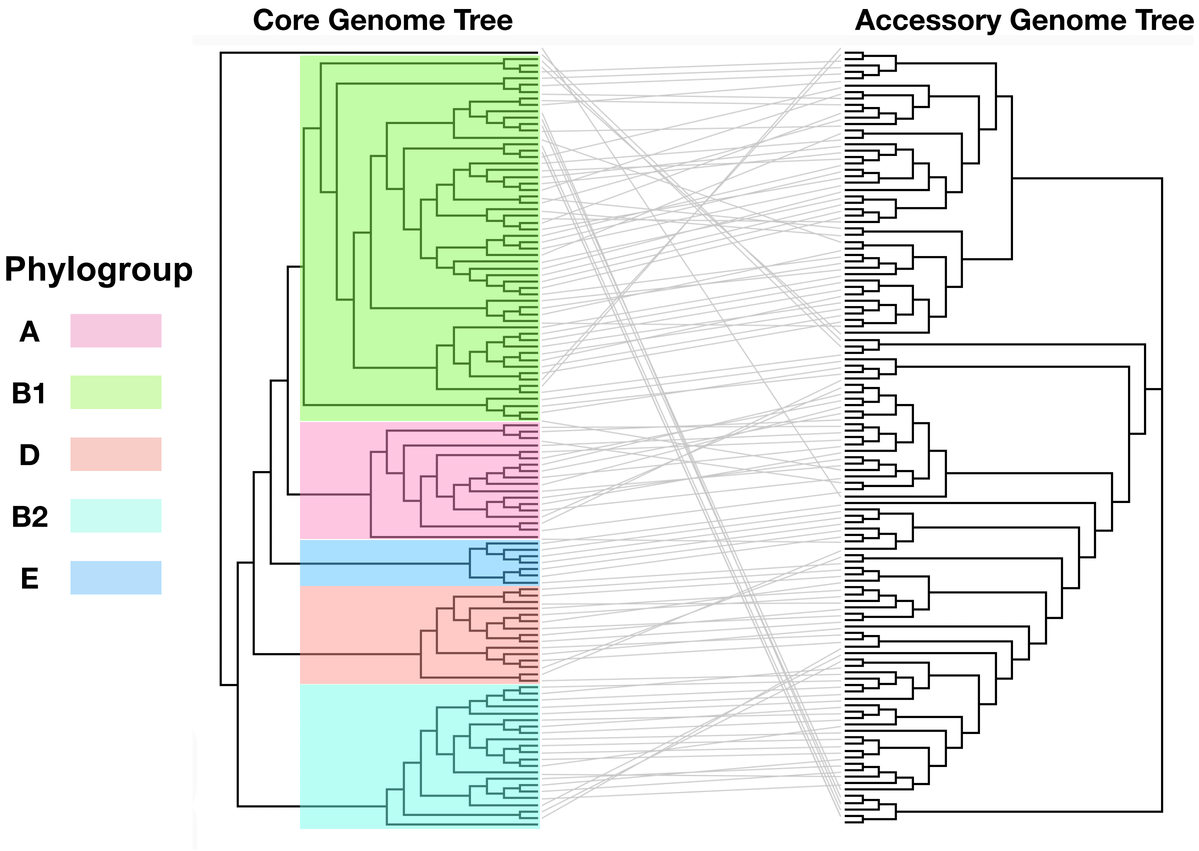


**Figure S2 The comparison between core genome and accessory genome trees**: The core genome and accessory genome trees were reconstructed from the SNPs in the core genome alignment and presence and absence genes pattern, respectively.

### Genomic epidemiology and evolution of *Escherichia coli* in wild animals: Supplemental Figures

Robert Murphy, Martin Palms, Ville Mustonen, Jonas Warringer, Anne Farewell, Danesh Moradigaravand, Leopold Parts

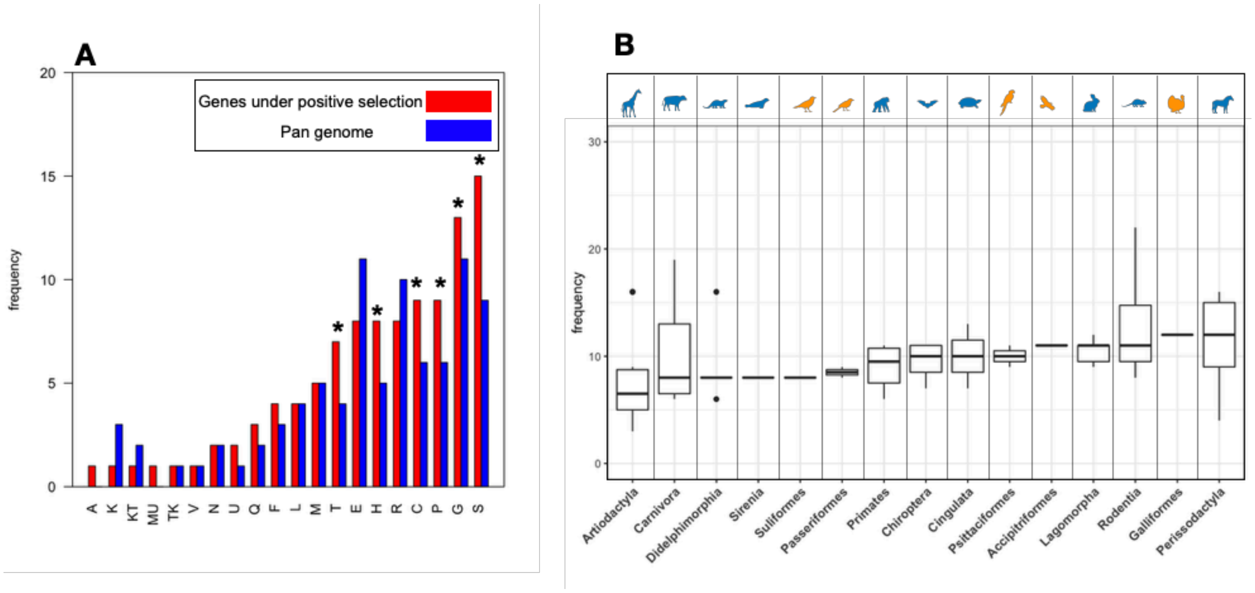


**Figure S2 Positive selection analysis:** A) The distribution of functional classifications for 256 genes that were under positive selection, compared with the baseline distribution in the pan genome. We excluded genes without assigned COG group. The S, G, P, C, H and T represent unknown functions, carbohydrate metabolism and transport, inorganic ion transport and metabolism, energy production and conversion, coenzyme metabolism and signal Transduction, respectively. The full interpretation for other classes may be found in (60). The asterisk sign shows 0.05 significance level from Student's test to assess the significance of the difference between groups. B) The distribution of genes under positive selection across taxonomic hosts orders.

**Genomic epidemiology and evolution of *Escherichia coli* in wild animals: Supplemental Figures**

Robert Murphy, Martin Palms, Ville Mustonen, Jonas Warringer, Anne Farewell, Danesh Moradigaravand, Leopold Parts

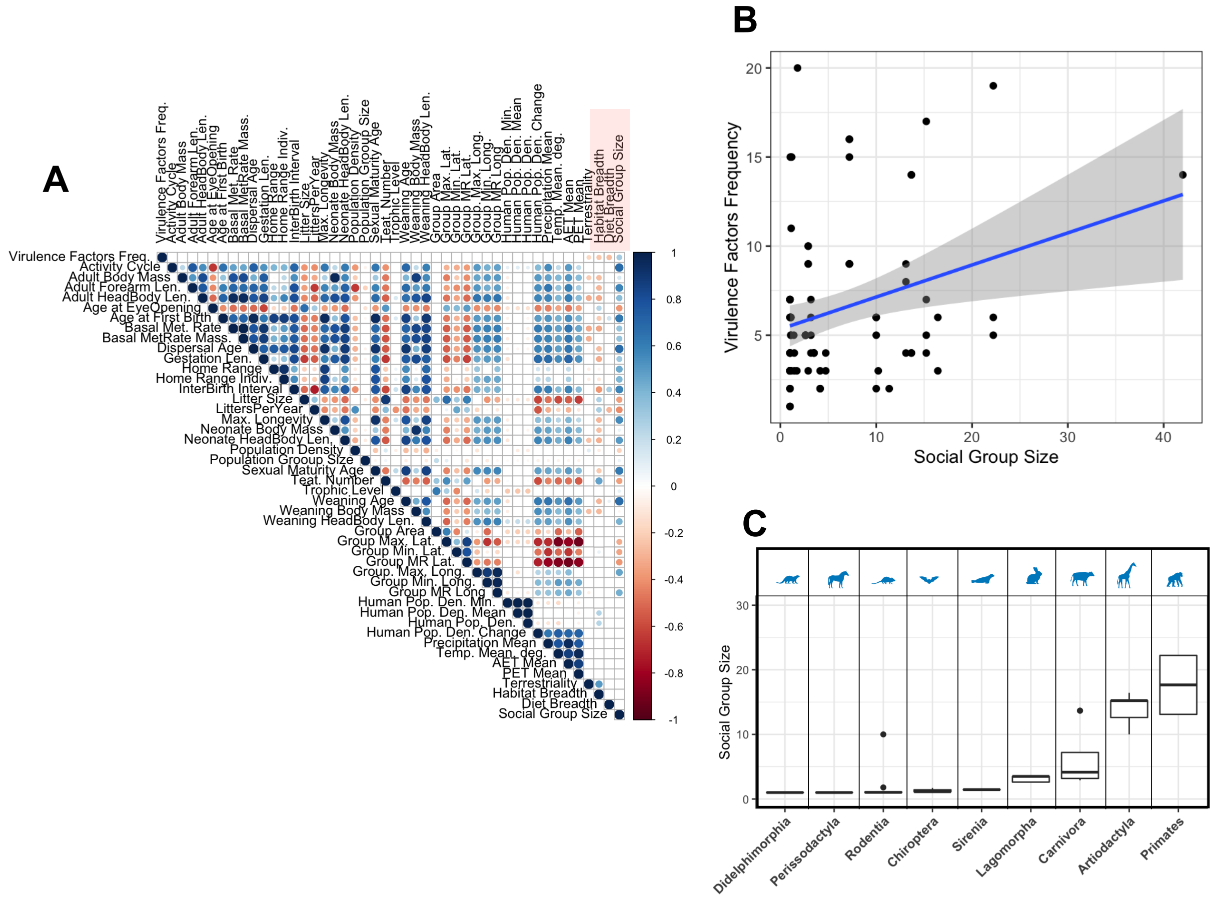


**Figure S4 Correlation between virulence factor counts and physiological and ecological attributes of wild hosts:** A) the correlation coefficient between the number of virulence genes and ecological features in the panTHERia dataset. The size and color of the circles correspond to the absolute value and direction of Spearman's rank correlation coefficient, respectively. Entries with insignificant correlation correlations, i.e. *p*-value< 0.01, are shown in blank. The red box shows the ecological features that are significantly correlated with virulence factor counts. B) The correlation between social group size and total number of virulence genes for the wild-host *E. coli* sample set. The blue line is the fitted linear regression model. The grey area corresponds to 95% confidence interval. The adjusted *R*-squared value is 0.27. C) Social group size values across host orders.

**Genomic epidemiology and evolution of *Escherichia coli* in wild animals: Supplemental Figures**

Robert Murphy, Martin Palms, Ville Mustonen, Jonas Warringer, Anne Farewell, Danesh Moradigaravand, Leopold Parts

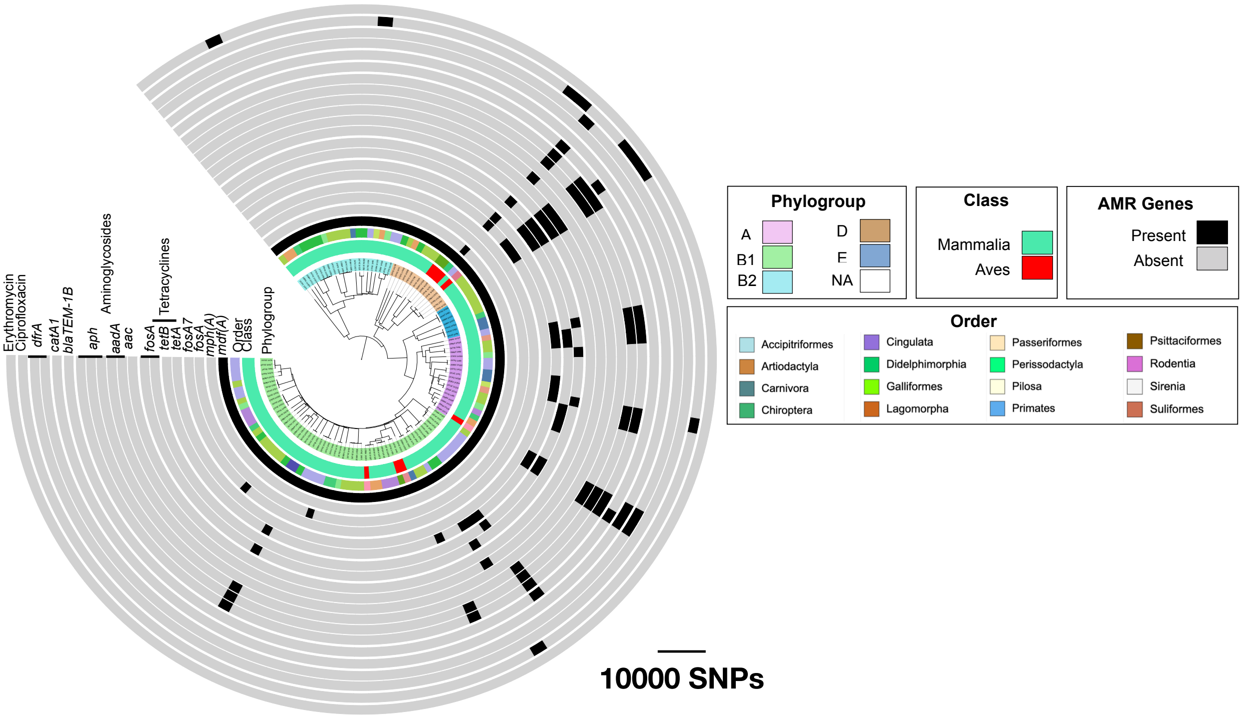


**Figure S5 The phylogenetic distribution of antimicrobial resistance genes.**: The tree was built from SNPs in the core genome, using the neighbour-joining method. For ciprofloxacin resistance, chromosomal mutations are collectively shown.
